# Consequences of combining sex-specific life-history traits

**DOI:** 10.1101/2020.01.03.892810

**Authors:** Vandana Revathi Venkateswaran, Olivia Roth, Chaitanya S. Gokhale

## Abstract

Males and females evolved distinct life-history strategies, reflected in diverse inter-linked life-history traits. The sex that allocates more resources towards offspring relies on an increased life span, and long life requires an efficient immune system. The other sex needs to attract mates and thus allocates its resources towards ornamentation, which may negatively correlate with investment into the immune defense. Such sex-specific resource allocation trade-offs are not always strictly female or male-specific but may depend on the overall resources allocated towards life-history traits. Informed by experimental data, we designed a theoretical framework that combines multiple life-history traits. We disentangled specific life-history strategies from particular sex, allowing us to include species with reversed sex-roles and male parental investment. We computed the lifetime reproductive success (combining fitness components from diverse sex-specific life-history traits) observing a strong bias in adult sex ratio depending on sex-specific resource allocation towards life-history traits. Overall, our work provides a generalized method to combine various life-history traits with sex-specific differences to calculate lifetime reproductive success. The results explain specific population-level empirical observations as a consequence of sexual dimorphism in life-history traits.

## Introduction

Reproductive success depends on the ability of individuals to produce offspring and survive. The lifetime reproductive success arises from trade-offs among various life-history traits. Theoretical and experimental studies have shown how multiple life-history traits define an individual’s lifetime reproductive success (Moore, 1990; Martin, 1992; Chapman and Partridge, 1996; Pusey et al., 1997; Fleming et al., 2000; Alonzo, 2002; Kalbe et al., 2009; Alonzo, 2010; Kelly and Alonzo, 2010). However, typically, these traits have been studied in isolation or the sex-specific differences were not accounted for. In this study, we present a model that addresses the interdependence of essential sex-specific life-history traits, aiming to obtain the lifetime reproductive success of both sexes. This approach sheds light on how these traits are contributing to an individual’s life-history. We present how various sex-specific traits affect an evolving population.

For reproduction, females contribute large, costly eggs and males small cheap sperm. This difference in gamete size is known as anisogamy (Bell, 1978). Likewise, the way resources are allocated towards various sexual and life-history traits differs between males and females. Here we focus on the sex-specific differences in three life-history traits, namely 1. Parental investment 2. Ornamentation and 3. Immunocompetence.

In many species, parental investment is not restricted to sperm and egg production. Parental investment (PI) is any behavioural and physiological investment by a parent provided to the offspring (Trivers, 1972, 2002). The sex that needs to allocate more resources towards the offspring strives for increased longevity since the parent that spends more time per reprodutive unit needs to survive longer to produce the same number of offspring when compared to the other sex that has lower time investment per reproductive unit. Increased longevity requires the allocation of resources into parasite defence and, hence, immunity (Roth et al., 2011; Lin et al., 2016). Intense, costly intrasexual competitions for obtaining mates are performed by allocating resources towards ornamentation (Hillgarth and Wingfield, 1997; Wong and Candolin, 2005; Andersson and Simmons, 2006). To this end, fewer resources may be available for the immune defence in the sex majorly investing in intrasexual interactions. This implies that both ornamentation and parental investment contribute to sexual immune dimorphism (Forbes, 2007; Nunn et al., 2008; Roth et al., 2011; Lin et al., 2016). Thus focusing only on one of these life-history traits in isolation will not shed light on the individual’s actual lifetime reproductive success; accounting for the interdependence between various traits is necessary.

We aimed for designing a framework in which multiple interlinked life-history traits can be studied simultaneously. Notably, we have constructed a holistic framework that captures sex-specific differences in parental investment, ornamentation and immune response and presents the consequences of the overall life-history of a sex. The two significant consequences that we observed are 1) skewed adult sex ratios and 2) different ratios of homozygous and heterozygous individuals between the sexes concerning immune alleles. We have discussed our findings in the light of empirical data from a broad range of animal taxa and diverse life-history traits.

## Model

We amalgamated approaches from standard population genetics and eco-evolutionary processes (Freeman and Herron, 2007; Otto and Day, 2007; Venkateswaran and Gokhale, 2019) (within and between populations) to investigate the interaction dynamics of multiple life-history traits (with sex-specific differences). We first developed a robust method (illustrated in Figure 1 to study the lifetime reproductive success (LRS) that arises from immune response, mating competition through ornaments and parental investment. Later, we used the LRS to investigate the consequences of combining the sex-specific traits that are part of an individual’s reproductive lifetime.

**Figure 1:**
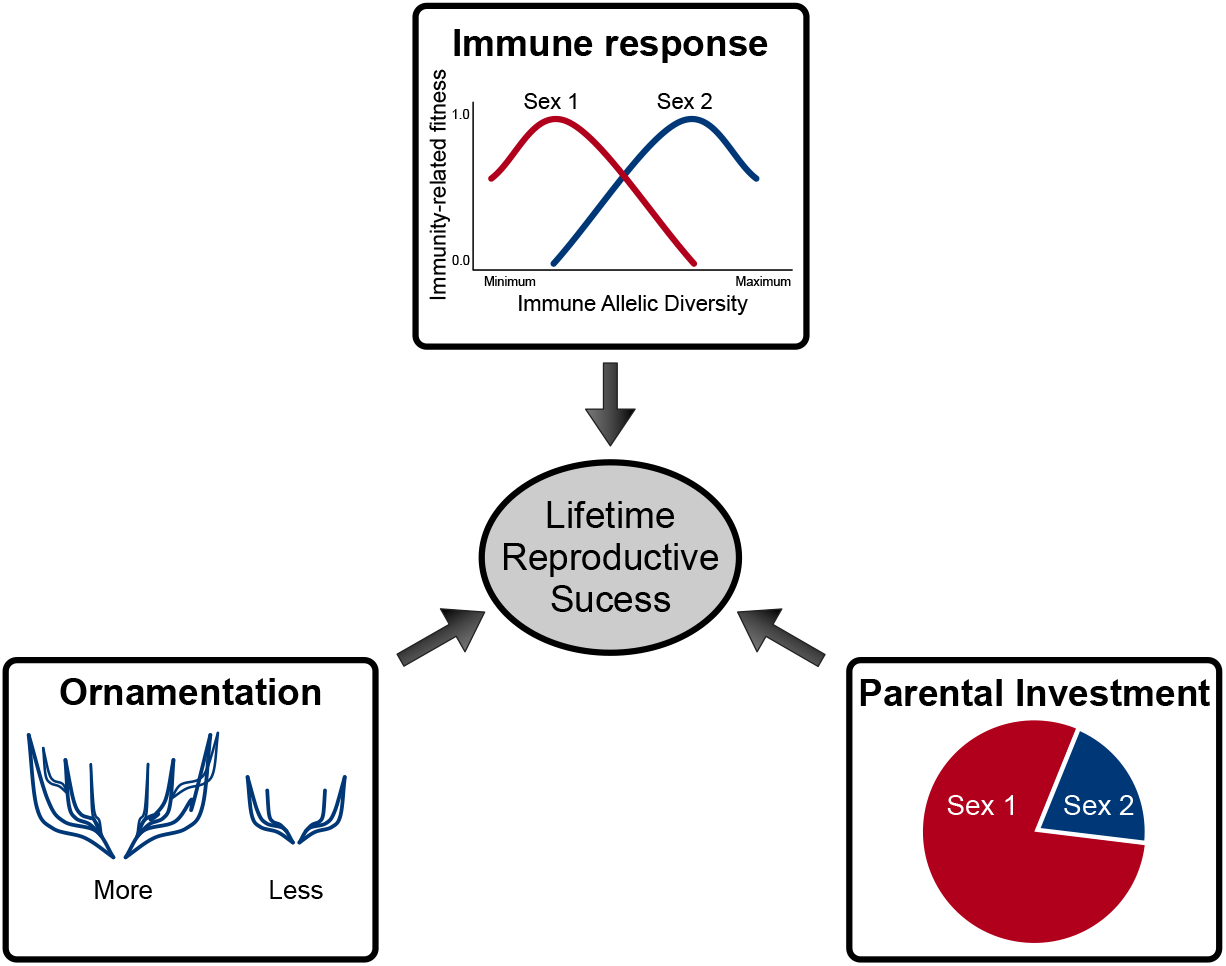
Model representation. Life-history traits affect the lifetime reproductive success. The fitness components from parental investment, immune system and ornamentation are offspring success, survival of the parent plus offspring and mating success, respectively. These contribute to an individual’s lifetime reproductive success. We assumed that Sex 1 provides more parental investment (PI) than Sex 2. The sex-specific fitness from parental investment is modeled as frequency dependent since the number of copulations in one sex depends on the availability of the other sex. The individuals of one sex also have different levels of ornamentation, which they use to attract individuals of the other sex as potential mates. The model uses evolutionary game theory which gives frequency dependent fitnesses of two types of individuals: those with more and those with lower level of ornaments. The individuals also differ in their immune genotypes. Each immune genotype yields a certain immunity-related fitness value that depends on the type and number of different immune alleles. The strength of immune response differs between sexes (sexual immune dimorphism). We modeled the evolution of these immune genotypes using population dynamics. Finally, the fitness obtained from parental investment, ornamentation and immune response were used to measure the lifetime reproductive success of an individual.

Consider the two sexes in a population, Sex 1 denoted by a filled circle •, and Sex 2 denoted by a diamond ◊. We first consider one autosomal immunity locus *A* having two alleles *A*_1_ and *A*_2_. The three distinct zygotes genotypes would be *A*_1_*A*_1_, *A*_1_*A*_2_ and *A*_2_*A*_2_. For Sex 1, which throughout this manuscript does major PI, the frequencies of the three genotypes are denoted by *x*_•1_, *x*_•2_, *x*_•3_. The fitnesses, of the same, are denoted by *W*_•1_, *W*_•2_ and *W*_•3_. Similarly, we denote the frequencies and fitnesses for Sex 2.

We used standard Mendelian segregation to model the evolution of the different types of individuals in the population. The genotype dynamics follow the segregation pattern (electronic supplementary material, ESM). As with normal Mendelian segregation, we assumed equal sex ratio; half of the offspring are Sex 1 and the other half, Sex 2.

### Lifetime reproductive success

The lifetime reproductive success of an individual, i.e. the overall fitness of an individual, is related to its immunocompetence (the ability to produce a healthy immune response following exposure to a pathogen), the ability to obtain mates, and its offspring success (Stoehr and Kokko, 2006; Kalbe et al., 2009; Kelly and Alonzo, 2010). Thus, in our model, the sex-specific fitness components resulting from immune response, ornamentation and parental investment give the lifetime reproductive success of individuals of a sex as shown in Figure 1. Below we introduce the fitness functions independently starting with immunity.

#### Immune response

A host’s immune allelic diversity helps eliminate a large number of pathogens and disease-causing agents. However, in some cases, having too high allelic diversity may reduce efficient immune responses, e.g. auto-immune diseases triggered by high Major Histocompatibility Complex (MHC) diversity. Thus, having an optimal number of immune allelic diversity (intermediate diversity) has been shown to be ideal in many systems (Nowak et al., 1992; Milinski, 2006; Woelfing et al., 2009). The host’s immune allelic diversity can be coarsely split into three parts: low diversity (*LD*, low efficiency of the immune system), intermediate or optimal diversity (*ID*, optimal immune efficiency), and high diversity (*HD*, might reduce the efficiency of the immune system). Recent experimental studies by Roved et al. (2017, 2018) and Jamie Winternitz and Tobias Lenz (personal communication) show that the optimal diversity could differ between the sexes. Based on these ideas, we have different cases that are shown in the Figure 2 for one immune locus *A* with two alleles *A*_1_ and *A*_2_ that gives three distinct parent and offspring genotypes *A*_1_*A*_1_, *A*_1_*A*_2_, and *A*_2_*A*_2_ denoted by *j* = {1, 2, 3}. We denote their immune responses by 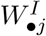 and 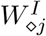 for genotypes *j* = {1, 2, 3} in the two sexes. In our model, we refer to immune allelic diversity as the number of different immune alleles in the immune loci. Later, a non-linear immune allelic diversity profile as shown in Figure 3, where the negative effect of *HD* is also addressed.

**Figure 2:**
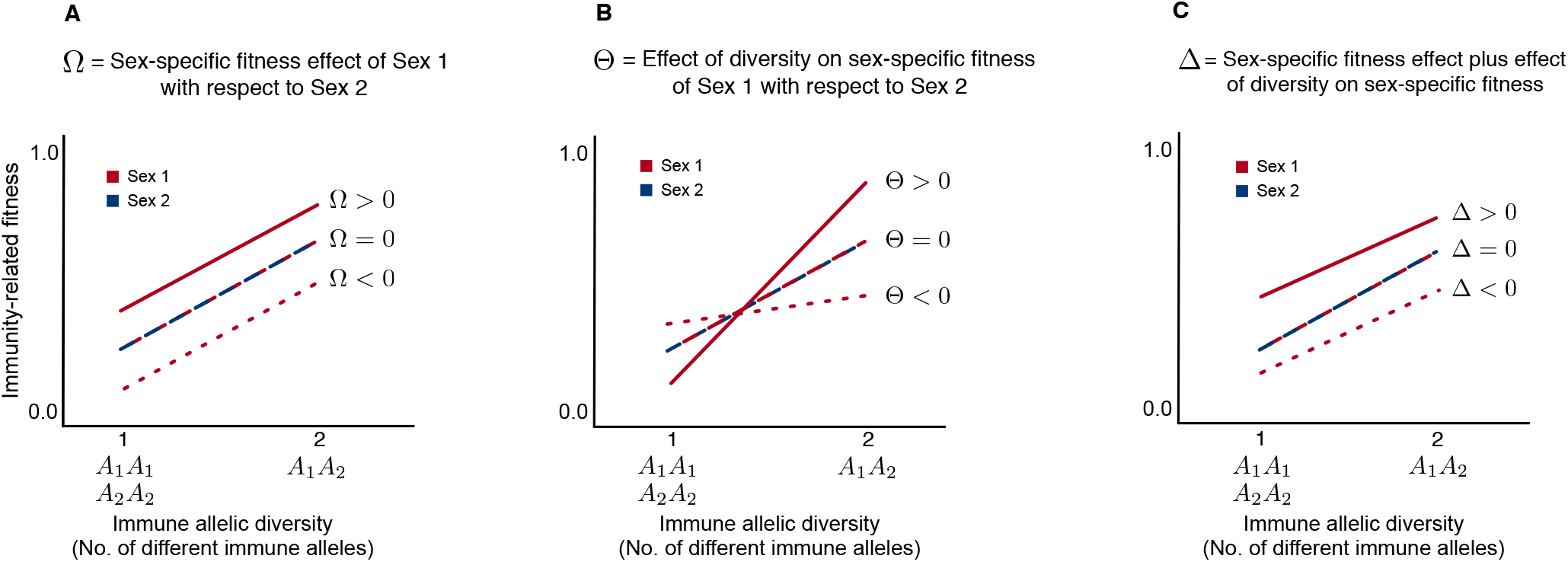
Schematic representation of different scenarios of sex-specific differences in host immunity-related fitness versus immune allelic diversity. We considered three distinct immune genotypes *A*_1_*A*_1_, *A*_1_*A*_2_, and *A*_2_*A*_2_ that result from mating between individuals having one immune gene locus *A* with two alleles *A*_1_ and *A*_2_ (Mendelian segregation, see ESM). Fitness positively correlates with the number of different alleles or allelic diversity (Apanius et al., 1997; Eizaguirre et al., 2009). So genotypes *A*_1_*A*_1_ and *A*_2_*A*_2_ (homozygotes) will have the same fitness value as they both have only one type of allele. But *A*_1_*A*_2_ (heterozygote) which has two different types of alleles will have a higher fitness. This is known as heterozygous advantage and occurs within both sexes. However, between the sexes, there can be sex-specific differences (Roved et al., 2017). This is shown in panels (A), (B) and (C). In (A), Ω > 0 would imply that Sex 1 will have a higher value of immune response as compared to Sex 2 for any given allelic diversity. When Ω < 0, Sex 1 has a lower values of immune response for any given allelic diversity as compared to Sex 2. Another situation is also possible: Sex 1 can have higher immune response for a homozygous locus, and lower immune response for a heterozygous locus when compared to Sex 2. This shown in (B), where Θ is the difference between the angles of the two lines. In (C), Δ differs from Ω by considering lines that are not parallel to each other i.e. case C is a combination of cases A and B. When both sexes have the same immune response patterns, Ω = Θ = Δ = 0.

**Figure 3:**
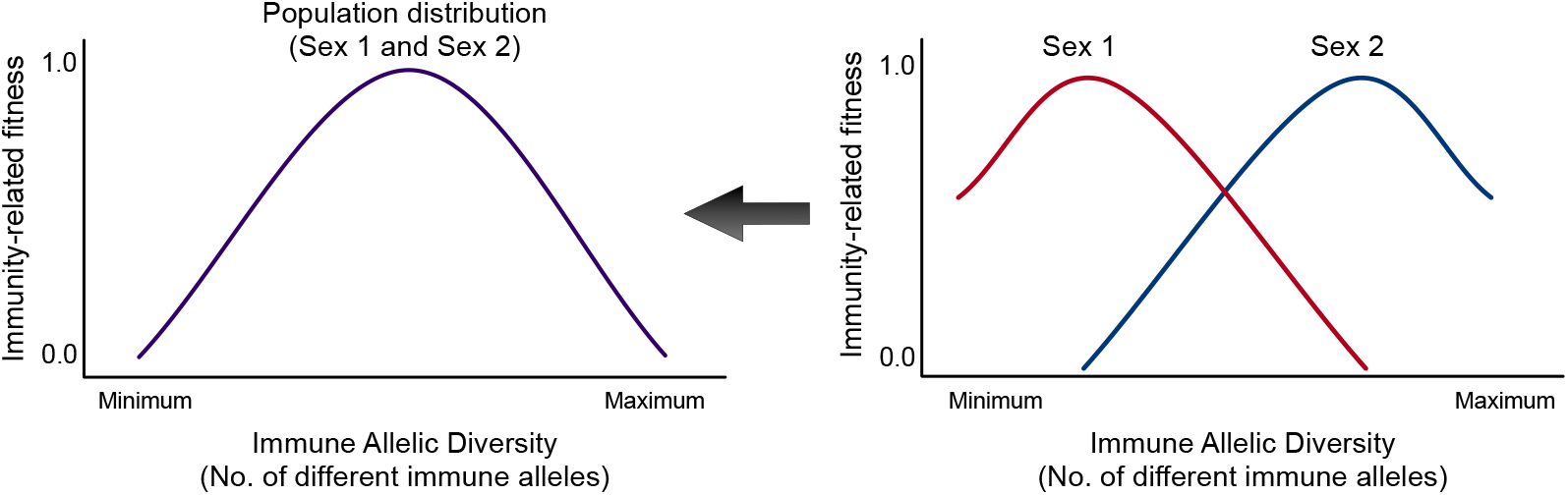
Schematic representation of host immunity-related fitness versus immune allelic diversity. For two immune gene loci *A* and *B* each having two alleles *A*_1_,*A*_2_ and *B*_1_,*B*_2_, there would be ten distinct zygote genotypes. The population will comprise of individuals with these genotypes. Their immune responses would depend on these genotypes. The probability of immune response might reduce if the individual has too many immunity allele diversity. In the case of MHC, the auto-immune effect of having high MHC allele diversity reduces the probability of immune response (Nowak et al., 1992; Milinski, 2006; Woelfing et al., 2009). Thus there is an optimal allele diversity, which gives the parabolic shape to the curve. Recent studies have shown that males and females can have different optimal diversities ((Roved et al., 2017, 2018) and Winternitz et al., unpublished). Plotted here are hypothetical sex-specific optima of immune allelic diversity (Roved et al., 2017). The realized population distribution is what is typically looked at, but in our study we consider sex-specific optima of immune allelic diversity. Some immune genes may follow completely different sex-specific patterns from the one shown here (Roved et al., 2017; De Lisle, 2019), and this model can be used for most kinds of immune genes.

These approaches can be generalised to any genetic system controlling the immune response or a completely different causal mechanism devoid of the genetic correlation. For example, the effect of nutrients and its effect of the immune system can be captured by a non-genetic model as well (Chandra, 1983). Thus, while we focus on the genetic mechanism in the current model, we stress that our framework is independent of the exact mechanism of how the immune response curves develop. Condition of an individual is directly proportional to immune response (resources allocated to self-maintenance, immune defense), which in turn determines survival (Stoehr and Kokko, 2006).

#### Parental investment

Both sexes pay the costs for initial PI, i.e. egg and sperm production (Hayward and Gillooly, 2011). Brooding the eggs is common in male and/or female birds, amphibians, replies, and fishes. Besides pregnancy by a single sex, one or both sexes of some species also exhibit external extended parental care, i.e. parental investment provided to the offspring after parturition or hatching (Trivers, 1972; Wade and Shuster, 2002; Trivers, 2002; Kokko and Jennions, 2003; Alonzo, 2010) (Figure 1). We assume that Sex 1 provides major PI (e.g. male sticklebacks, male pipefish, most female mammals). The fitness from PI will depend on the relative abundance of the other sex and are given by, 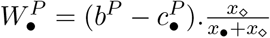 and 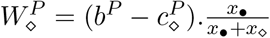. Here, *b*^*P*^ is the benefit (offspring produced) from PI while 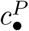 and 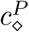 are the costs for PI by Sex 1 and Sex 2, respectively. The frequency of Sex 1 equals *x*_•_ = *x*_•1_ + *x*_•2_ + *x*_•3_ and the frequency of individuals in Sex 2 equals *x*_⋄_ = *x*_⋄1_ + *x*_⋄2_ + *x*_⋄3_. Since we have assumed that Sex 1 provides maximum parental investment, 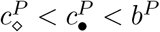.

#### Ornamentation

Mating competitions occur among individuals of the same-sex to attract and obtain mates. This competition is performed through fights, nuptial gifts, nests, sexual signals, ornament display and various types of ‘attractiveness’. We refer to all of these as ‘ornaments’. Ornamentation is a costly signal (Zahavi, 1977; Andersson and Simmons, 2006; Milinski, 2006; Kurtz, 2007)); but the investment into ornaments is beneficial as ornamentation will in most cases increase the chances of acquiring mates (Carranza et al., 1990; Petrie et al., 1991; Berglund et al., 1997; Wong and Candolin, 2005).

When Sex 2 participates in mating competition as shown in Figure 1, two types of Sex 2 individuals were considered in this interaction: one type displays more ornaments (*MO*) and the other type displays less ornaments (*LO*). Members of Sex 2 that win this competition game (based on their and their opponents’ strategies) are chosen by members of Sex 1. Therefore Sex 2 consists of six types of individuals - *x*_*⋄j,MO*_ and *x*_*⋄j,LO*_ where the genotype *j* = {1, 2, 3}. The frequency-dependent fitnesses from these interactions are as 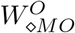 and 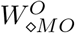 (see ESM for details).

### Overall dynamics

The lifetime reproductive success is a multiplicative effect of the fitness arising from immune response, ornamentation and parental investment (Stoehr and Kokko, 2006; Kelly and Alonzo, 2010) as shown in the ESM. Using the LRS values in the Mendelian population dynamics, we can obtain the combined interaction dynamics of each type of individuals in the population (details and calculations in the ESM). The population is divided into nine types of individuals - the three genotypes (*j*) of Sex 1, *x*_•*j*_, and the three genotypes of Sex 2 further split according to ornamentation into *x*_*⋄j,MO*_ and *x*_*⋄j,LO*_. We refer to them as simply *x*_*i*_ with *i* as the type of individual. The classical selection equation from population genetics (Crow and Kimura, 1970) gives the evolution of the frequency *x*_*i*_ having average fitness *W*_*i*_ (Crow and Kimura, 1970; Schuster and Sigmund, 1983; Hofbauer and Sigmund, 1998; Gokhale et al., 2014). The equation can be written as,

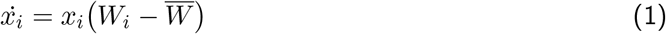

where 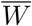 is the average population fitness.

## Results

### Linear immune allelic diversity profile: single locus

The diversity levels in the immune alleles can result in differing immune response (e.g. MHC homozygotes and heterozygotes, are known to have different immune responses (Apanius et al., 1997; Eizaguirre et al., 2009)). For one immune locus with two alleles, higher allele diversity induces the specificity of the immune response as shown Figure 2. The negative effect of very high diversity is not considered here. Beyond the null model of immune allelic diversity, we include different cases of sexual conflict (Roved et al., 2018) (Figure 2).

When we assume that both sexes are not involved in mating competition, i.e. ornamentation competition game is neutral; we can vary the cost of PI and the immune response curves (shown in Figure 2). The resulting equilibrium frequencies are shown in Figure 4. When the cost of PI is zero, and there is no sex-biased difference in immune response, we observe that the sex ratio is 1 : 1. Here, we focus on the adult sex ratio (ASR) (Kokko and Jennions, 2008). The classical definition of ASR is “number of males:population size”. However, in this model, Sex 1 could be male or female. We thus here define the term ASR as the ratio between Sex 1 and Sex 2. Since in every generation offspring are produced in equal sex ratios (see ESM), what we obtain is the sex ratio of the offspring after they become adults, perform mating interactions and parental investments. The frequency of Sex 1 decreases with increasing PI. However, Sex 1 increases in frequency under certain cases of sexual conflict over the immune allelic diversity (see Δ > 0, Ω > 0, or Θ ≠ 0 in Figure 2). The results after including mating competitions are plotted in the figures in the ESM.

**Figure 4:**
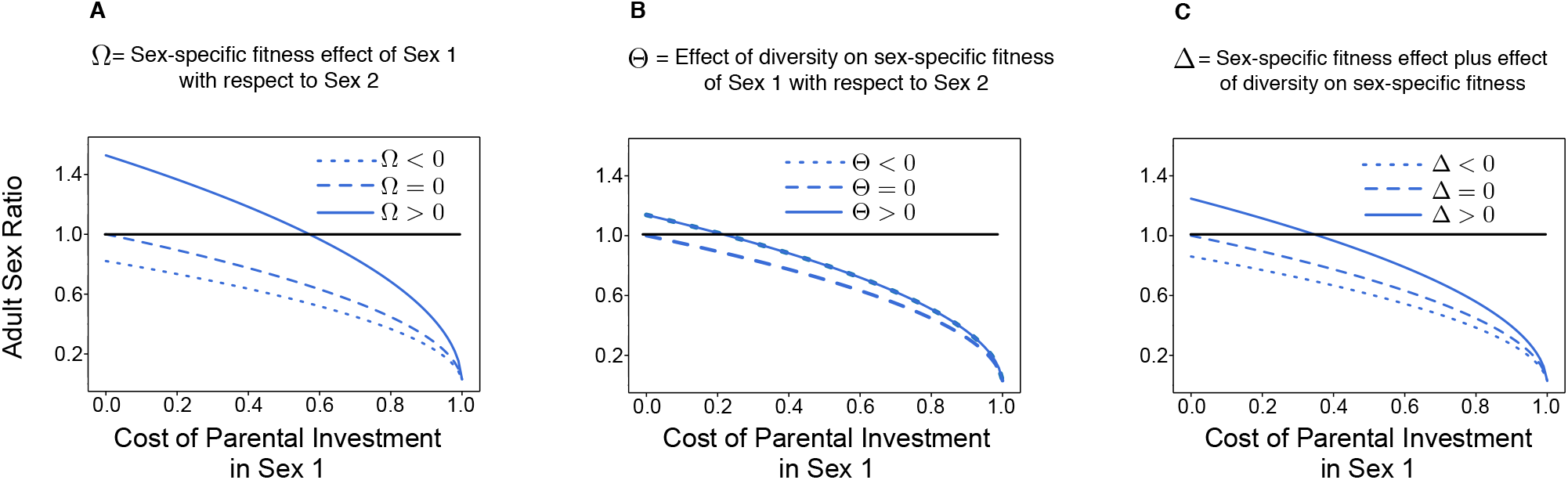
Adult sex ratio (Sex 1: Sex 2) for varying parental investment (PI) and various cases of sexual conflict within immune allelic diversity as shown in Figure 2. The ornamentation game is neutral, i.e. no selection acting on it (details in the ESM). As maintained throughout this study, Sex 1 does maximum PI. Sex 2 does negligible PI. Therefore, its cost is set to zero, i.e. 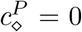. The black line highlights the even adult sex ratio, i.e. 1:1. In (A), (B) and (C): When the cost of PI = 0 and there is no sex difference in immune response (Ω = Δ = Θ = 0), the obtained adult sex ratio is 1:1. In (A) and (C): when PI increases, the frequency of Sex 1 drops as PI is costly. When Ω > 0 and Δ > 0, this sex difference in immune response compensates for the cost of PI. The fall in frequency of Sex 1 is lower than when Ω = 0 and Δ = 0 and Sex 1 has higher frequency than Sex 2 for most values of PI cost. However, when Ω < 0 and Δ < 0, Sex 1’s frequency decreases with an increase in PI. In (B): Frequency of Sex 1 is lower than Sex 2 for most values of PI cost for most Θ values. Moreover, Θ < 0 and Θ > 0 give the same results. The above results highlight the fact that sexual conflict within immune allelic diversity can increase (when Ω > 0 and Δ > 0) or reduce (when Ω < 0, *δ* < 0, almost all Θ) the adult sex ratio.

Under selection, the obtained genotypes deviated from the Hardy-Weinberg equilibrium (see Figures S.3, S.5 and S.4 of the ESM). One sex has a higher number of heterozygotes when compared to the other sex. In this setup, the heterozygous immune genotype (*A*_1_*A*_2_) has a higher immune response than the homozygous genotypes *A*_1_*A*_1_ and *A*_2_*A*_2_ (Figure 3). Thus, an increase in heterozygotes within one sex compared to the other would also mean that this sex has a higher average activity of the immune system. A recent study with wild songbird populations, where the number of heterozygotes and homozygotes, even under selection, turned out to be equal between the sexes (Roved, 2019). However, this could be the result of a particular immune response profile, parental investment and ornamentation costs in that species. Different profiles of sexual conflict within the immune allelic diversity would determine different ratios of homozygotes and heterozygotes. More empirical studies with various model organisms should shed light on how species show diverse ways of sexual conflict within the immune allelic diversity.

### Nonlinear diversity profile

In a multi-loci scenario, one can include non-linear density profiles (Nowak et al., 1992; Woelfing et al., 2009) as shown in Figure 3. Across species, different sex-specific immune response profiles can be found, depending on the sex-specific selection and phenotypic divergence (Uekert et al., 2006; Love et al., 2008; Oertelt-Prigione, 2012). We hypothesize two such scenarios,

- the optimal diversity of immune alleles for both sexes is the same, but the immune responses at this optimal diversity could differ between the sexes (for instance, females are more prone to acquiring autoimmune diseases; sex hormones such as estrogen, testosterone also affect immune response (Hillgarth and Wingfield, 1997; Törnwall et al., 1999; Whitacre, 2001) or,
- the two sexes have different optimal diversity of immune alleles, and the immune response at this optimal diversity is the same for both sexes. For instance, as shown in Roved et al. (2017, 2018), males and females have a different optimal diversity, where males need a higher allelic diversity to mount a maximum immune response. We considered such a scenario for this study (see Figure 3).

As done for the one locus scenario, we assume that only the number of different alleles, i.e. allele diversity produces unequal fitness.

### Adult sex ratio in various species

Our results showed that a sexual conflict within immune allelic diversity and varying parental investment might result in adult sex ratio bias. The effect of ornamentation also plays an essential role in skewing adult sex ratios, as shown in Figure 5. Diverse reproducing species have distinct ornamentation and parental investment costs. Figure 5 shows the values of adult sex ratios that our model predicts for a wide range of species.

**Figure 5:**
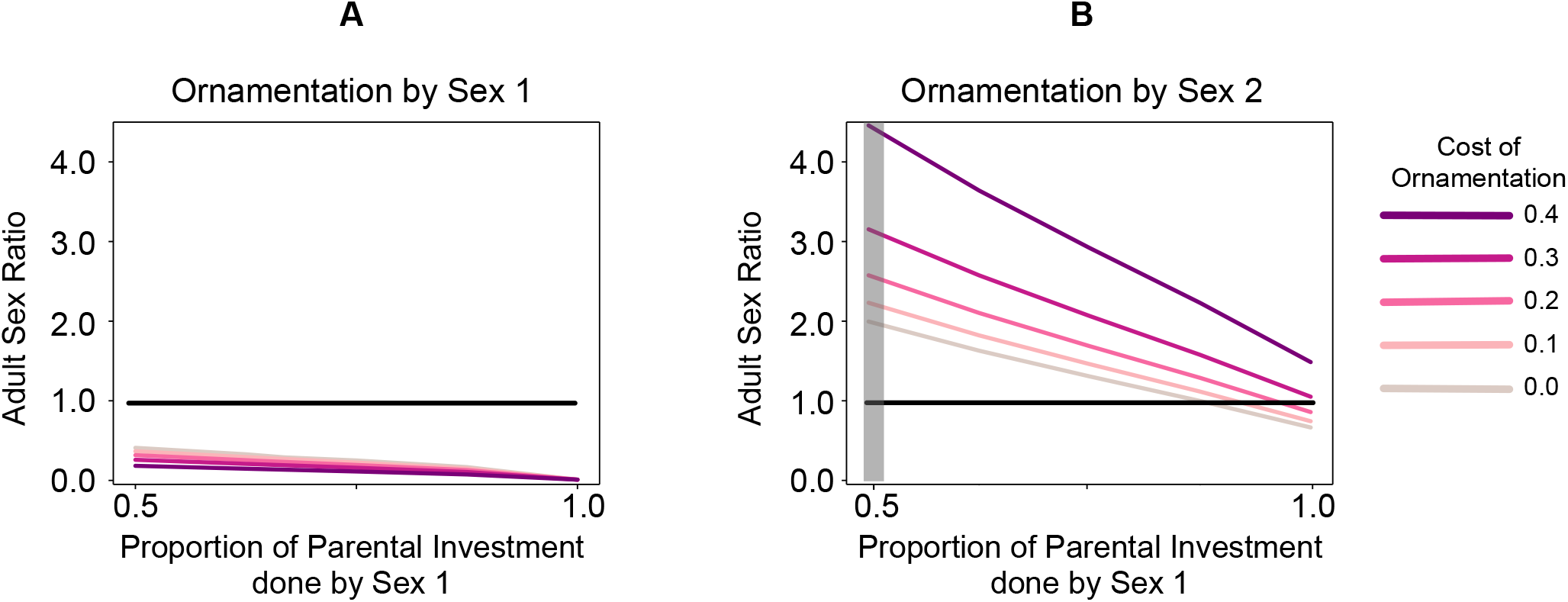
Qualitative difference in the adult sex ratio for diverse polygamous species with varying parental investment (PI) and ornamentation costs. As defined throughout the manuscript, Sex 1 is the major PI provider. For these calculations, we used the sexual conflict Case 4 shown in Figure 3. (A) Species such as sticklebacks where one sex performs both ornamentation and most PI. We observe that frequency of Sex 1 descends as its PI cost increases and this further decreases with a rise in its ornamentation cost. (B) The panel highlighted in gray shows bi-parental investment scenarios. In species where Sex 1 does most PI and Sex 2 performs elaborate mating competitions, the frequency of Sex 1 reduces with increasing PI. However, this value grows with ascending ornamentation cost in Sex 2. Note that for certain ornamentation and PI values, the adult sex ratios are equal. As shown by previous studies on multiple interactions between traits (Venkateswaran and Gokhale, 2019), even in the case where the cost of ornamentation is equal to zero in the mating competition game, the mere presence of that game will deviate the frequency of Sex 2 from a scenario where there is no ornamentation game.

## Discussion

Various life-history traits during the reproductive lifespan of an individual determine its lifetime reproductive success (Stoehr and Kokko, 2006; Kalbe et al., 2009; Kelly and Alonzo, 2010). We have presented a model framework where a combination of several individual life-history traits can be studied simultaneously. As shown in many empirical studies, the ASR has an impact on sex-specific differences and roles (Liker et al., 2013; Székely et al., 2014; Liker et al., 2015; Henshaw et al., 2019). Our model predicted that the interaction of sex-specific life-history traits results in a biased adult sex ratio (ASR) (Pipoly et al., 2015). We showed that the vice versa is also possible (Kokko and Jennions, 2008), i.e., our results showed that ASR is a consequence of sex-specific differences. Our model incorporates the fact that fitness is a complex entity (Doebeli et al., 2017). The overall lifetime reproductive success is a combination of fitness values arising from individual life-history traits. The precise individual-level variation in traits has population-level consequences, i.e. a skew in adult sex ratio (see Figures S.1 and S.2 in the ESM). Here, the females and males of one generation mate and produce equal numbers of daughters and sons in the next generation. Therefore, at birth, the sex ratio of every generation was 1:1. The life-history traits are passed on from parents to offspring. Thus, even though every generation starts with an equal sex ratio, their sex-specific traits change the adult sex ratio in every generation until it reaches an equilibrium state.

If a sex invests more in ornamentation and maximum parental investment (e.g. stickleback males), the ASR will be biased towards the sex that bears negligible costs for the same traits (e.g. female sticklebacks) (Hagen and Gilbertson, 1973)). Thus, the high costs for contributing to both PI and ornamentation cannot be compensated (Daly, 1978) (Figure 5.A).

In birds and free-spawning fish, both sexes exhibit similar levels of parental investment (equally little parental investment by both sexes in case of free-spawning fish) (Perrone Jr and Zaret, 1979; Gross and Sargent, 1985; Cockburn, 2006). Our model shows that these species could exhibit equal ASR for particular parental investment and ornamentation levels (see Figure 5.B). However, in species where males have a higher ornamentation level, the ASR will be biased. For instance, free-spawning species such as the Atlantic salmon where males have elaborate ornaments, show a highly skewed adult sex ratio (7:1 ratio of males to females) (Mobley et al., 2019). Therefore, the high sex ratio values are shown in the grey shaded region of Figure 5.B matches what we can find in nature.

When one sex makes the maximum parental investment, while the other displays ornaments, ASR is biased towards the sex that invests more in parental care (Figure 5.B). Consider the pipefish species *N. ophidion* where males glue the eggs on the belly and thus perform partial parental investment (Berglund et al., 1986). In contrast to pipefish species with placenta-like structure and an active transfer of nutrients and oxygen to the embryo (e.g. *S. typhle* (Berglund et al., 1986; Smith and Wootton, 1999)), *N. ophidion* only provide partial parental investment. We thus expect a decrease in frequency of *S. typhle* males compared to *N. ophidion* males (Berglund and Rosenqvist, 2003). However, with increasing ornamentation in females, the frequency of males increases. Ornaments are costly as they make the bearer more vulnerable to predation. According to Bateman’s principle (Bateman, 1948), the reproductive success of the sex that performs mating competition depends on the number of mating events. The sex-limited by parental investment will have to live longer for more reproductive events to achieve the same reproductive success as the males (Roth et al., 2011). Thus sex differences in parental investment, ornamentation and immunity (Trivers, 1972; Hedrick and Temeles, 1989; Trivers, 2002; Roved et al., 2017) may also give rise to sexual differences in longevity, an important life-history trait (Austad, 2006; May, 2007).

Our model can be used to determine the lifetime reproductive success using fitness arising from sex-specific differences in life-history traits of a particular sex. Studying the combined dynamics of life-history traits highlights population-level consequences such as skewed adult sex ratio (Trivers, 2002; Kokko and Jennions, 2008) emerging due to sex-specific differences in life-history traits. With the aid of more empirical work directed towards investigating sexual conflict within the immune allelic diversity and other life-history traits, we can obtain a more in-depth understanding of the overall life-history of a sex and the species in general. Disruptive selection leads to sexual dimorphism and in models that use tools like adaptive dynamics, traits that go through evolutionary branching may end up as two sex-specific traits, i.e. sexual dimorphism. Recent studies addressed how coevolution of traits and resource competition drive the evolution of sexual dimorphism (Bolnick and Doebeli, 2003; Stoehr and Kokko, 2006; Vasconcelos and Rueffler, in press). Work by Vasconcelos and Rueffler (in press) demonstrated that even weak trade-offs between life-history traits could result in evolutionary branching that leads to the coexistence of the types. In this study, we investigated the eco-evolutionary consequences of interplay between two or more sex-specific life-history traits. Along with empirical evidence that matches our qualitative predictions, our model suggests a skewed adult sex-ratio.

The functions in our model that describe fitness from parental investment and ornamentation consider polygamous species. While many sexually reproducing animals are polygamous, species like seahorses are monogamous throughout their lifetime (Vincent and Sadler, 1995). The trade-offs between ornamentation, parental investment and immunocompetence in monogamous species would be different. For instance, they may not have to bear the costs of attracting mates after one brooding season. Our model can be modified to study the effect of integrating monogamous mating patterns. Concerning immune genes such as the ones of the MHC, genetically dissimilar individuals mate more often as the evolutionary incentive is to produce optimal MHC diversity offspring (Milinski, 2006; Woelfing et al., 2009; Kalbe et al., 2009; Eizaguirre et al., 2009). To this end, mating is not random. Aspects of a model by Kirkpatrick (Kirkpatrick, 1982) for two autosomal loci with a female mating preference for a trait that occurs in males is a potential extension of our model. Finally, novel studies directed at sexual conflict within the MHC and other immune genes as done by Roved et al. (2018) shall be very beneficial in providing further knowledge of how sex-specific immune defences manifest in different systems with distinct sex-specific ornamentation and parental investment patterns.

## Electronic Supplementary material

### S.1 One locus with two alleles: Separate population into males and females

If the population is separated into the two sexes, Sex 1 which could be male (or female) denoted by a solid circle symbol •, and Sex 2 which could be female (or male) denoted by a diamond symbol ⋄. We stick to calling the sexes as Sex 1 and Sex 2 instead of males and females (and we also do not use the standard ♀ and ♂ symbols as it might be misleading) because we want to show a generalized idea of the dependence of sexual immune dimorphism on the amount of parental investment (or mating competition and other factors) given and not to the sex itself.

For Sex 1, let frequency of *A*_1_*A*_1_ = *x*_•1_, frequency of *A*_1_*A*_2_ = *x*_•2_ and frequency of *A*_2_*A*_2_ = *x*_•3_. Similarly, for Sex 2, let frequency of *A*_1_*A*_1_ = *x*_⋄1_, frequency of *A*_1_*A*_2_ = *x*_⋄2_ and frequency of *A*_2_*A*_2_ = *x*_⋄3_.

In Sex 1, let the fitness of individuals with genotype *A*_1_*A*_1_ = *W*_•1_, fitness of *A*_1_*A*_2_ = *W*_•2_ and fitness of *A*_2_*A*_2_ = *W*_•3_. Similarly, for Sex 2, let the fitness of individuals with genotype *A*_1_*A*_1_ = *W*_⋄1_, fitness of *A*_1_*A*_2_ = *W*_⋄2_ and fitness of *A*_2_*A*_2_ = *W*_⋄3_. The sex that performs mating competitions (say, Sex 2) is further divided into individuals with Less or More Ornamentation (*LO* or *MO*). Through Mendelian population dynamics we obtain the of frequency of each genotype at subsequent generations (Gokhale et al., 2014),

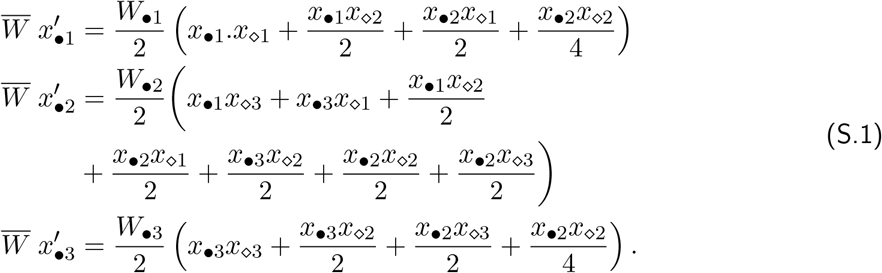

and

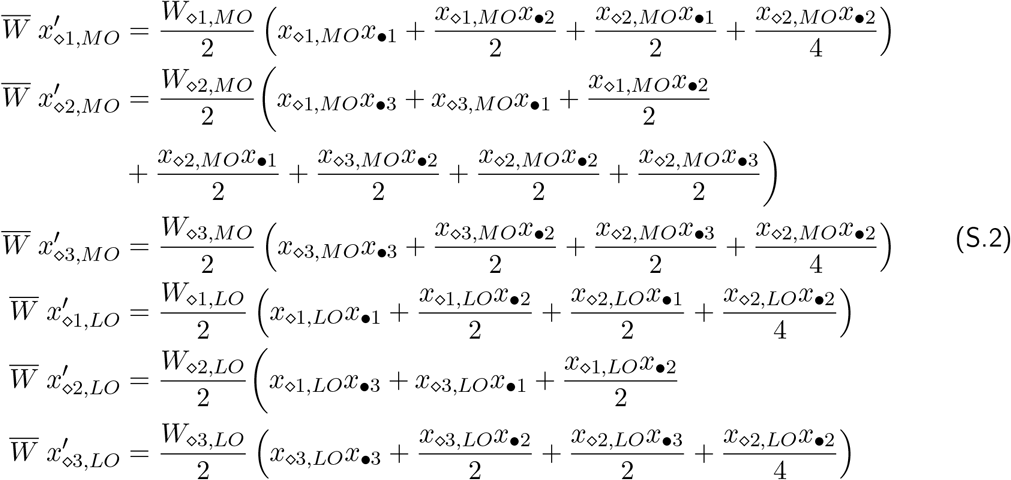

where 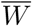 is the average fitness of all genotypes. Also, 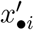 and 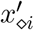 is the change in frequencies of the genotypes *i* (for the different sexes) with time. Also, here we assume equal sex ratio; half of the offspring are males and the other half, females.

Now, let *W*_⋄1_ = *W*_•1_ = *W*_1_, *W*_⋄2_ = *W*_•2_ = *W*_2_ and *W*_⋄3_ = *W*_•3_ = *W*_3_. where *W*_*⋄i*_ = *W*_*⋄i,MO*_ + *W*_*⋄i,LO*_. Then,

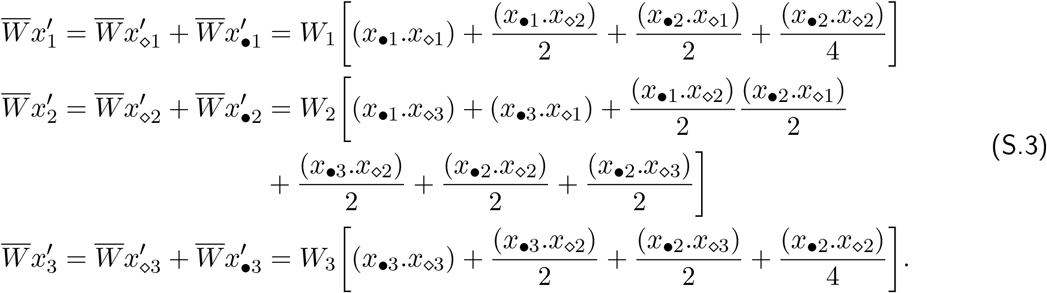

### S.2 Mating competition

Mating competition is performed through fights, sexual signals, nuptial gifts, ornament display and various types of attractiveness. We shall refer to all of these as ‘ornaments’. Let us assume there are individuals of two types in this interaction: ones that display more ornaments (MO) and ones that display less (LO). Consider the mating competition interaction between individuals of Sex 2. For the three different genotypes *i* the population in Sex 2 will consist of six different kinds of individuals, *x*_*⋄j,MO*_ and *x*_*⋄j,LO*_.

We model this interaction as an evolutionary game (Maynard Smith, 1986; Sigmund and Nowak, 1999). The payoff matrix is written as,

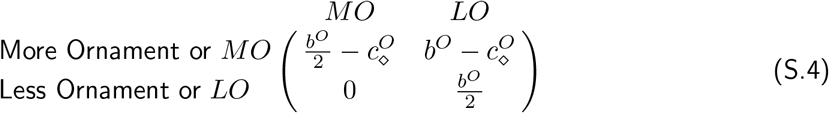

where *b*^*O*^ is the benefit arising from mating competitions, i.e. mating gain and 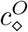 is the cost that Sex 2 bears to maintain ornament(s). The frequency dependent fitnesses resulting from these interactions are given by,

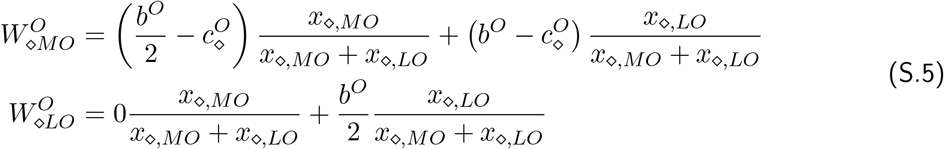

where 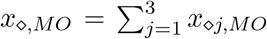 and 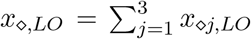. Sexual selection consists of two parts: mating competition and mate choice. While members of Sex 2 compete with each other in this ‘mating game’, the members of Sex 1 also chose their partners. Having *MO* strategy is ideal for attracting mates. Therefore, if the condition, 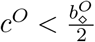 is taken into account in this payoff matrix, it ensures that members of Sex 2 that win this competition game (based on their and their opponents’ strategies) are chosen by members of Sex 1.

The payoff matrix (S.4) is an interaction between a pair of individuals, i.e. two player game. We can extend this to *d*-players (Gokhale and Traulsen, 2014; Chen et al., 2017) and the payoffs are given by,

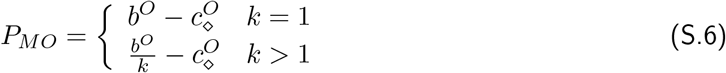

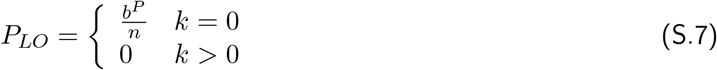

where *k* is the number of *MO* (More Ornament) players and *n* is the total number of players. *k* and *n* can vary between the sexes.

### S.3 One locus: Dynamics

**Figure S.1:**
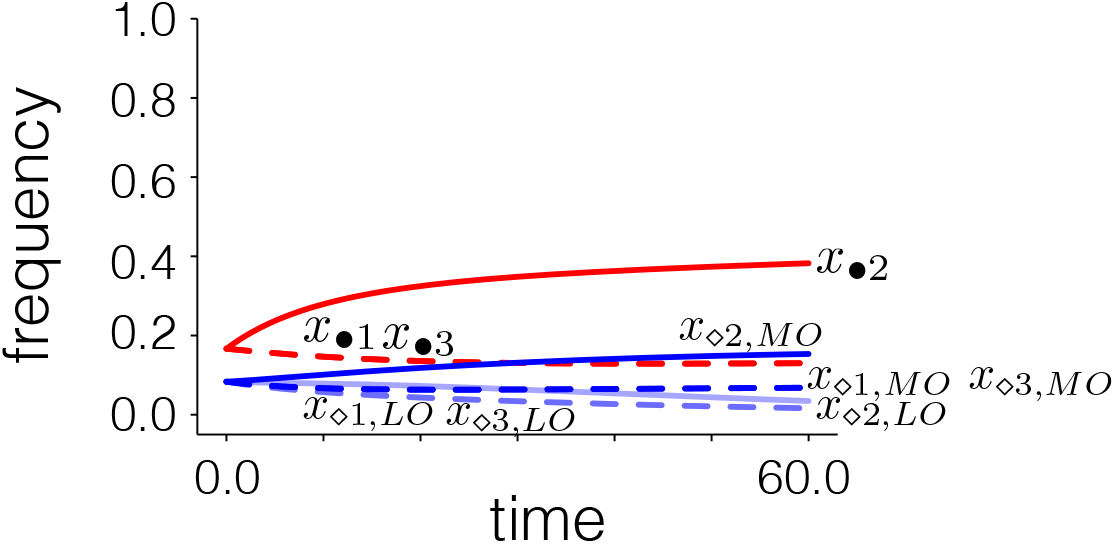
Evolution of frequency of all possible type of individuals in a population that exhibits sexual dimorphism in immunity and ornamentation, and sex difference in parental investment. When sex 1 performs major parental investment and individuals of Sex 2 perform mating competitions, then the population will have nine types of individuals - *x*_•*i*_,*x*_*⋄i,MO*_ and *x*_*⋄i,LO*_ for the three immunity genotypes *i*. For the results shown in this figure, 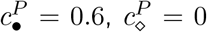 and 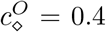. Fitness from immune response comes from Ω > 0 of the linear immune allelic diversity vs immune response profile in Figure 2 in the main article. Red lines are for Sex 1 and blue for Sex 2. The solid lines are for the heterozygous genotype and dashed lines for the homozygotes. The lighter blue lines in Sex 2 are for individuals with low ornamentation.

If we consider that sex 1, undergoing significant parental investment does not involve in mating competitions, and individuals of sex 2 perform mating competitions, then the population will have nine types of individuals - *x*_*•i*_,*x*_*⋄i,MO*_ and *x*_*⋄i,LO*_ for the three genotypes *i*. We shall refer to them as *x*_1_, *x*_2_, *x*_3_, *x*_4_, *x*_5_, *x*_6_, *x*_7_, *x*_8_ and *x*_9_.

The lifetime reproductive success of each type within a sex is a multiplicative combination of mating gains, fertility and survival probability (Stoehr and Kokko, 2006; Kelly and Alonzo, 2010). These are given by,

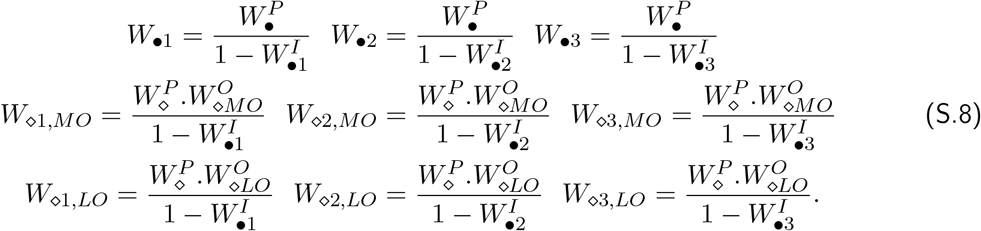

Here, 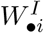 and 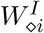 are the fitness from immune responses (survivability) of type *i* for Sex 1 and Sex 2 as described in the main text. Similarly, 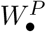 and 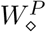 are the fitness that arise from parental investments performed by members of Sex 1 and Sex 2, respectively. The fitness from More and Less ornamentation (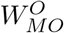 and 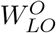 are as defined in the previous section. Using equations (S.1) and (S.2), we can obtain the average fitnesses for each type of individuals in the population. For Sex 1 they are given by,

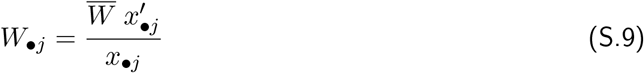

where *j* = {1, 2, 3} and 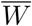 is the average fitness of all types. For Sex 2 they are given by,

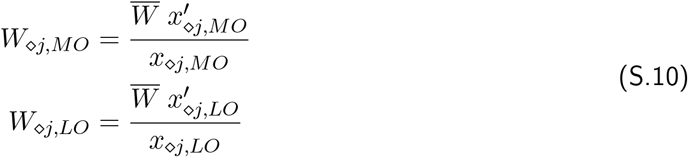

where again *j* = {1, 2, 3}. Here, *MO* and *LO* correspond to individuals with more and less ornamentation, respectively. From equations (S.9) and (S.9) we know that there are nine different types of individuals whose frequencies can be just described by *x*_*i*_ for *i* = {1, 2, 3, …9} and their respective average fitnesses are denoted by *W*_*i*_ (for *i* = {1, 2, 3, …9}).

Using the above given equations we have,

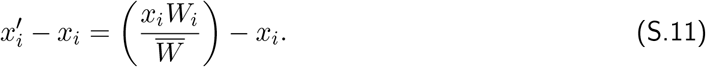

The classical selection equation (Crow and Kimura, 1970; Hofbauer and Sigmund, 1998) that gives the evolution of each type (see Figure S.1) is then obtained by taking the time derivative of (S.11) given by,

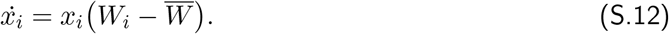

The frequencies of all types reach an equilibrium value at some time point. The equilibrium is used in the results throughout this ESM and the main article.

The frequency of each sex is a summation of frequencies of all types of individuals in a sex. Figure S.2 shows how the frequency of the sexes changes with sex-specific differences in immuno-competence, parental investment, and ornamentation.

**Figure S.2:**
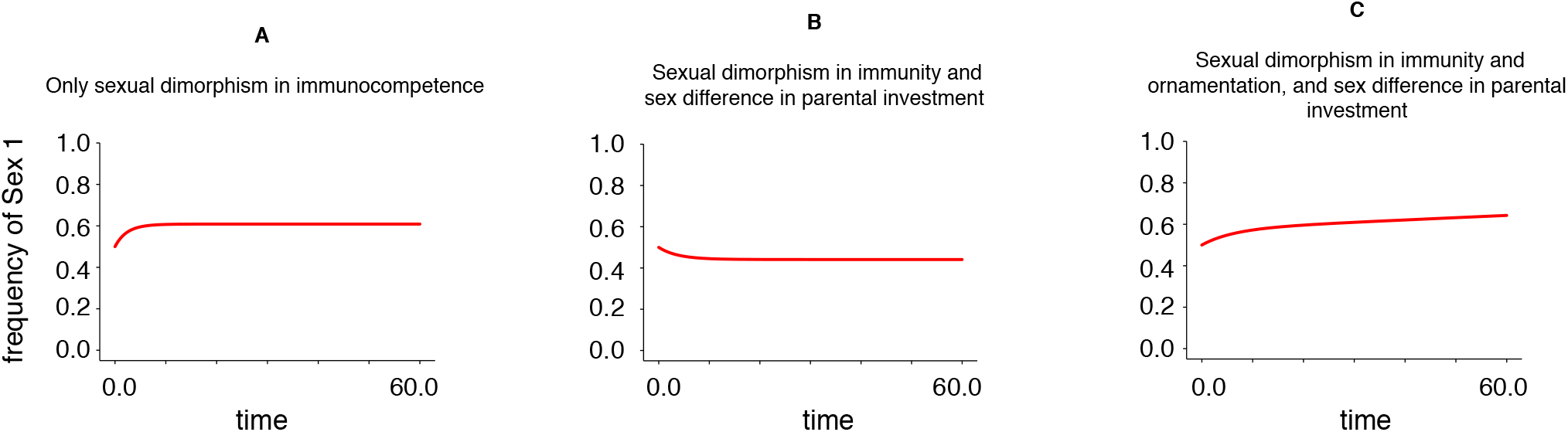
Evolution of frequency of Sex 1 in the population. Since frequency of Sex 1 (*x*_•_) and frequency of Sex 2 (*x*_⋄_) equals unity, *x* equals 1 − *x*_•_. These frequencies are obtained by summing up all types of individuals within the sexes. (A) When Sex 1 has a higher values of immune response a compared to Sex 2 for all immune allelic diversity (Ω). (B) When condition A is met, but Sex 1 also performs parental investment, while Sex 2 does not. (C) When conditions A and B are met, and Sex 2 also exhibits ornamentation. The sex-specific traits evolve over generation (time) by selection and therefore, get passed on to subsequent generations (for example, case C is shown in Figure S.1). Therefore, even when the sex ratio is kept equal among offspring at every generation, their sex-specific characteristics change their frequency in the population.

### S.4 One locus: Results

#### Heterozygosity vs Homozygosity

Under Hardy Weinberg or when all traits are neutral, the number of heterozygous and homozygous individuals within a sex would be equal. However, under selection (through different probabilities of immune response for homozygotes and heterozygotes), varying cost of parental investment and ornamentation the number of heterozygotes and homozygotes would deviate from neutrality. An increase in heterozygotes within one sex compared to the other would also mean than it has a higher immune response on average. When we allow for selection to act on all the three factors (parental investment, immunity genes and ornamentation), we can observe their combined effect on the increase in frequency of heterozygous individuals within a sex (results shown in Figures S.3,S.4 and S.5).

### S.5 Two loci having two alleles each

#### Population dynamics with separation of population into males and females

For Sex 1, let the frequency of *A*_1_*B*_1_|*A*_1_*B*_1_ = *f*_•_(*A*_1_*B*_1_|*A*_1_*B*_1_) = *x*_•1_, *f*_•_(*A*_1_*B*_1_|*A*_1_*B*_2_) = *x*_•2_, *f*_•_(*A*_1_*B*_2_|*A*_1_*B*_2_) = *x*_•3_, *f*_•_(*A*_1_*B*_1_|*A*_2_*B*_1_) = *x*_•4_, *f*_•_(*A*_1_*B*_2_|*A*_2_*B*_1_) = *x*_•5_, *f*_•_(*A*_1_*B*_2_|*A*_2_*B*_2_) = *x*_•6_, *f*_•_(*A*_2_*B*_1_|*A*_2_*B*_1_) = *x*_•7_, *f*_•_(*A*_2_*B*_1_|*A*_2_*B*_2_) = *x*_•8_, *f*_•_(*A*_2_*B*_2_|*A*_2_*B*_2_) = *x*_•9_ and *f*_•_(*A*_1_*B*_1_|*A*_2_*B*_2_) = *x*_•10_. Similarly, for Sex 2.

From Mendelian population dynamics (as done in the one locus case), the frequency of the homozygotes in Sex 1 will be:

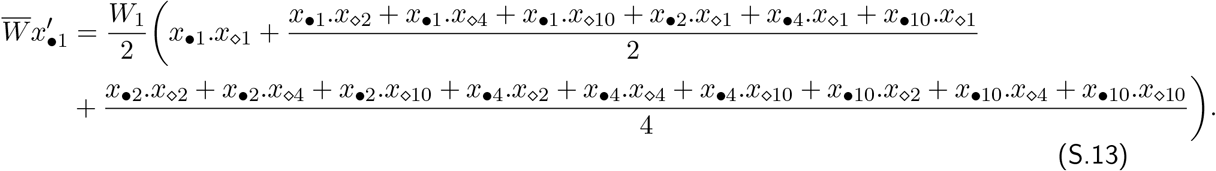

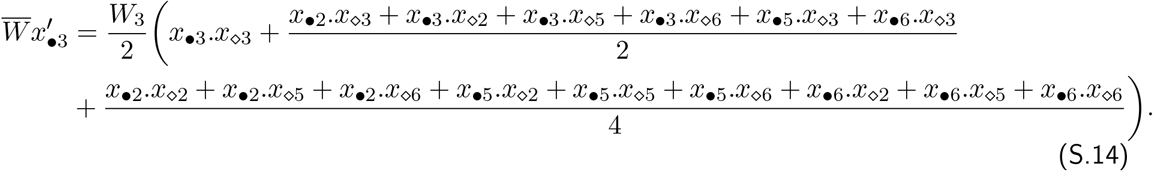

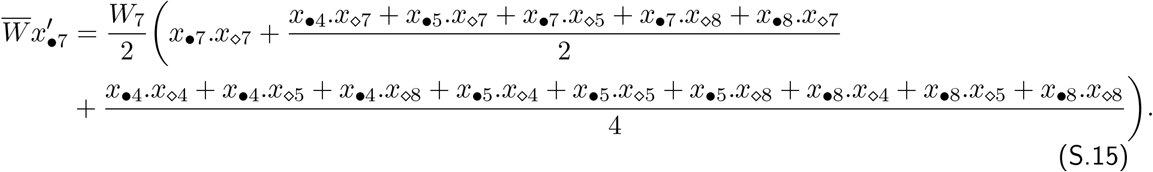

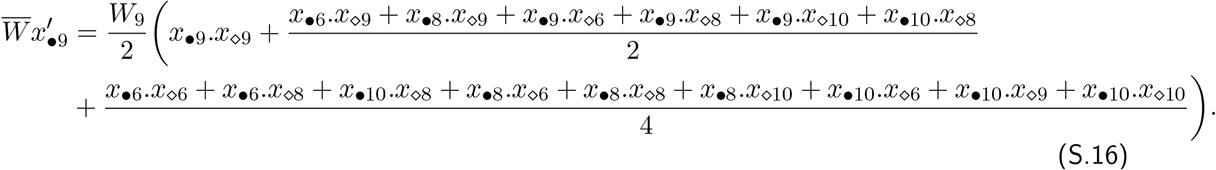

Frequency of the single heterozygotes will be:

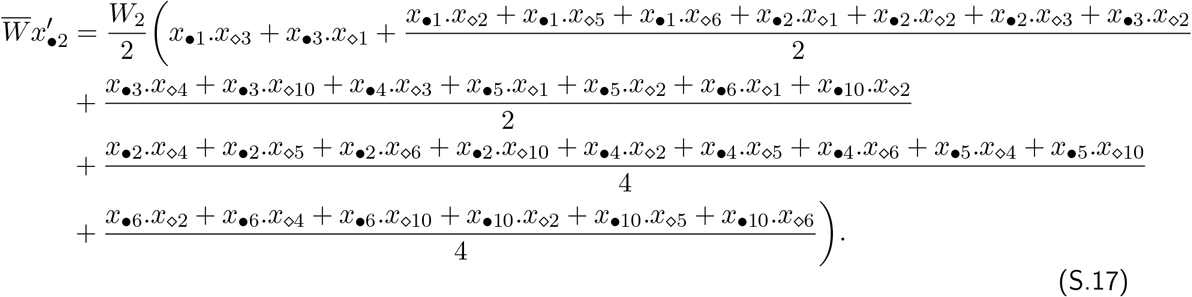

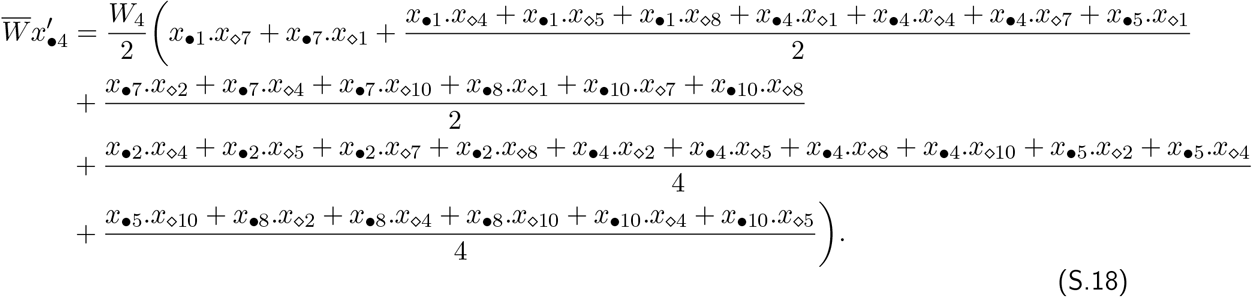

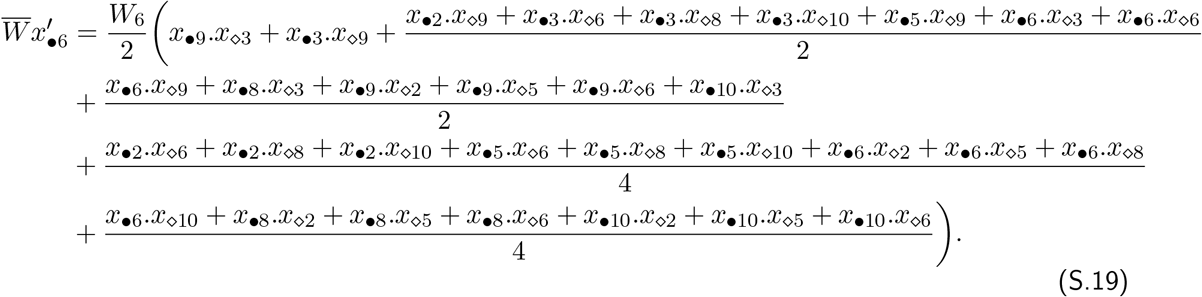

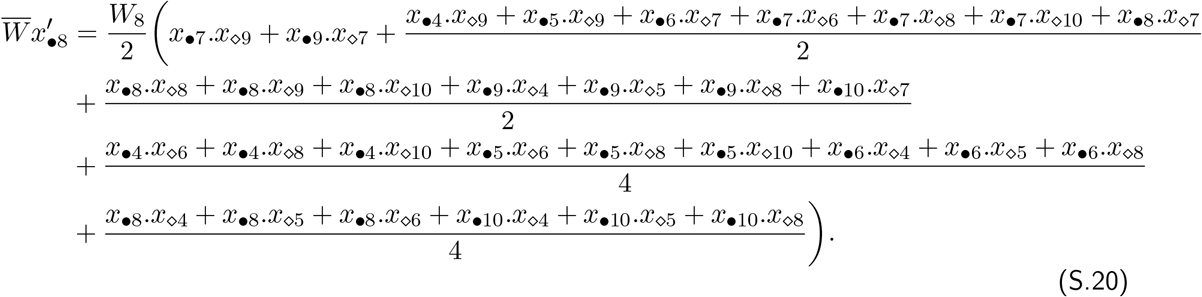

Frequency of the double heterozygotes will be:

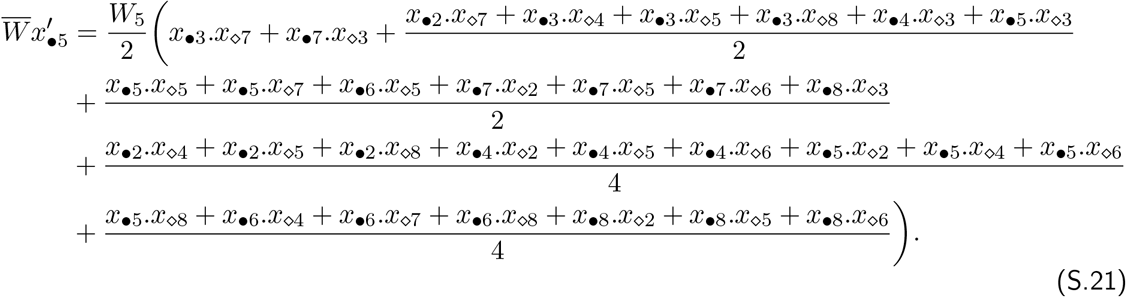

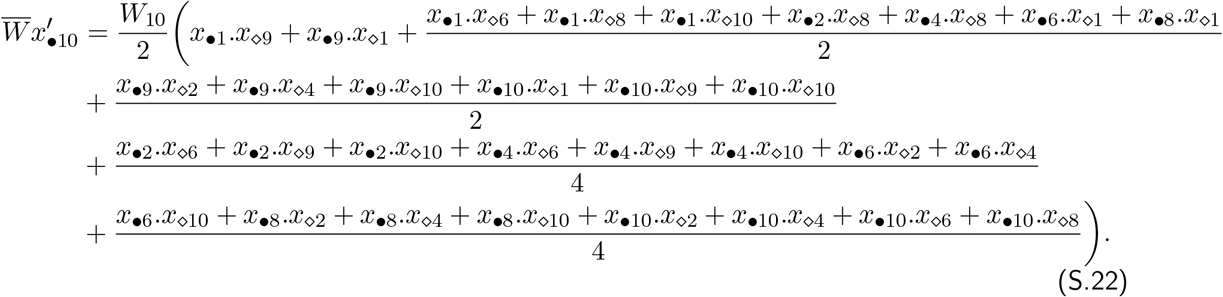

Here, the *W*_*i*_s are the fitnesses of each genotype *i* with frequency *x*_*i*_ and 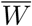 is their mean fitness. Similarly, we can obtain the frequencies of the genotypes in Sex 2.

**Figure S.3:**
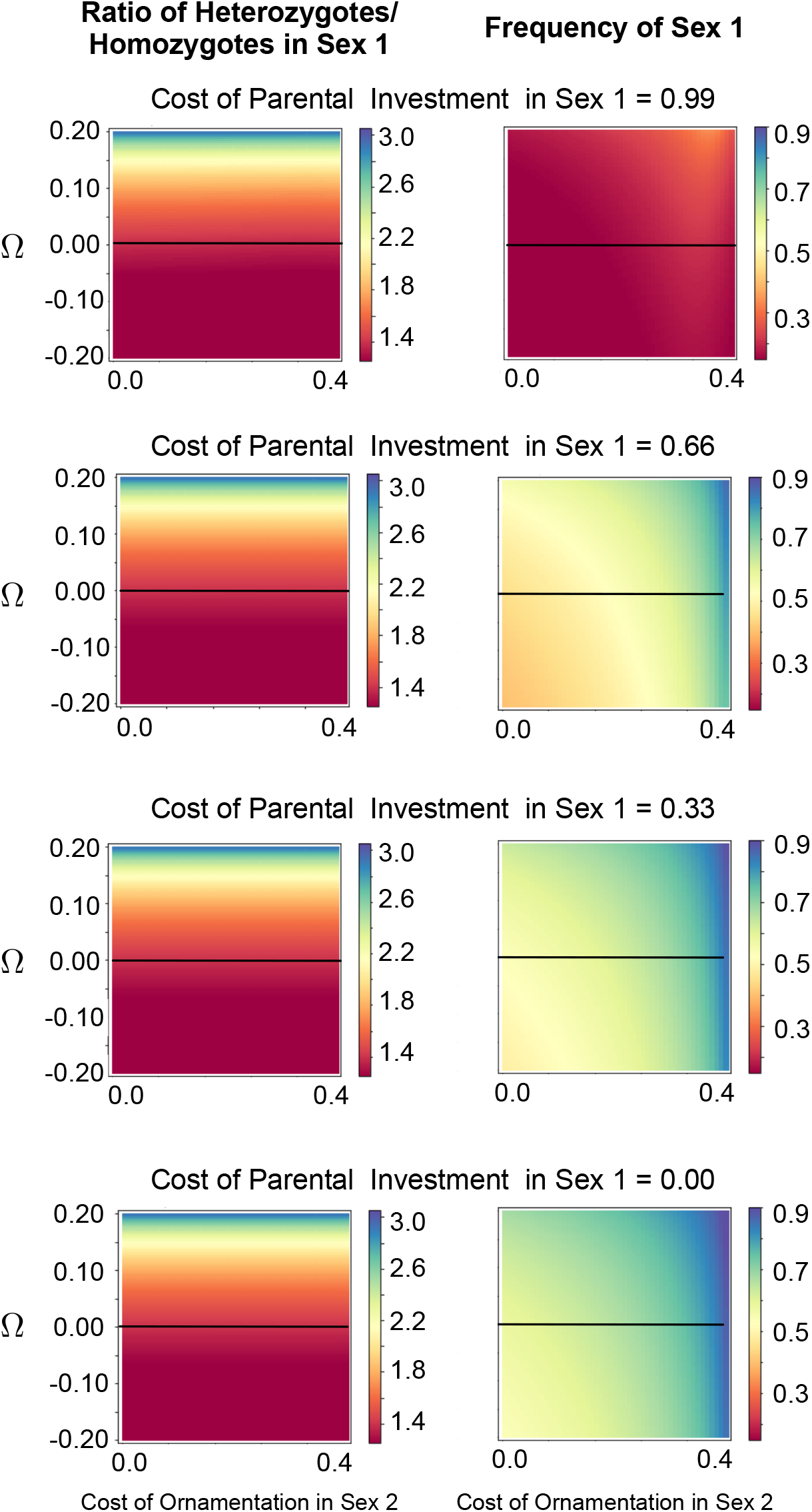
Ratio of Heterozygotes: Homozygotes in Sex 1 and the frequency of Sex 2 for a full range of Ω. The parameter Ω is a measure of the sex difference in immune response through sexual conflict within the MHC as shown in Figure 2. in the main article. It represents the sex-specific fitness effect of Sex 1 relative to Sex 2. When Ω = 0, there is so sex-specific difference in immune response. There is no effect of ornamentation and parental investment (PI) on the ratio of allele diversity. However, Ω has an effect on this ratio. All factors: coat of PI, cost of ornamentation and Ω have an effect on the frequency of the sexes. Thought the effect of Ω is not profound, the cost of ornamentation in Sex 2 and cost of PI in Sex 1 reduce their frequency, respectively.

**Figure S.4:**
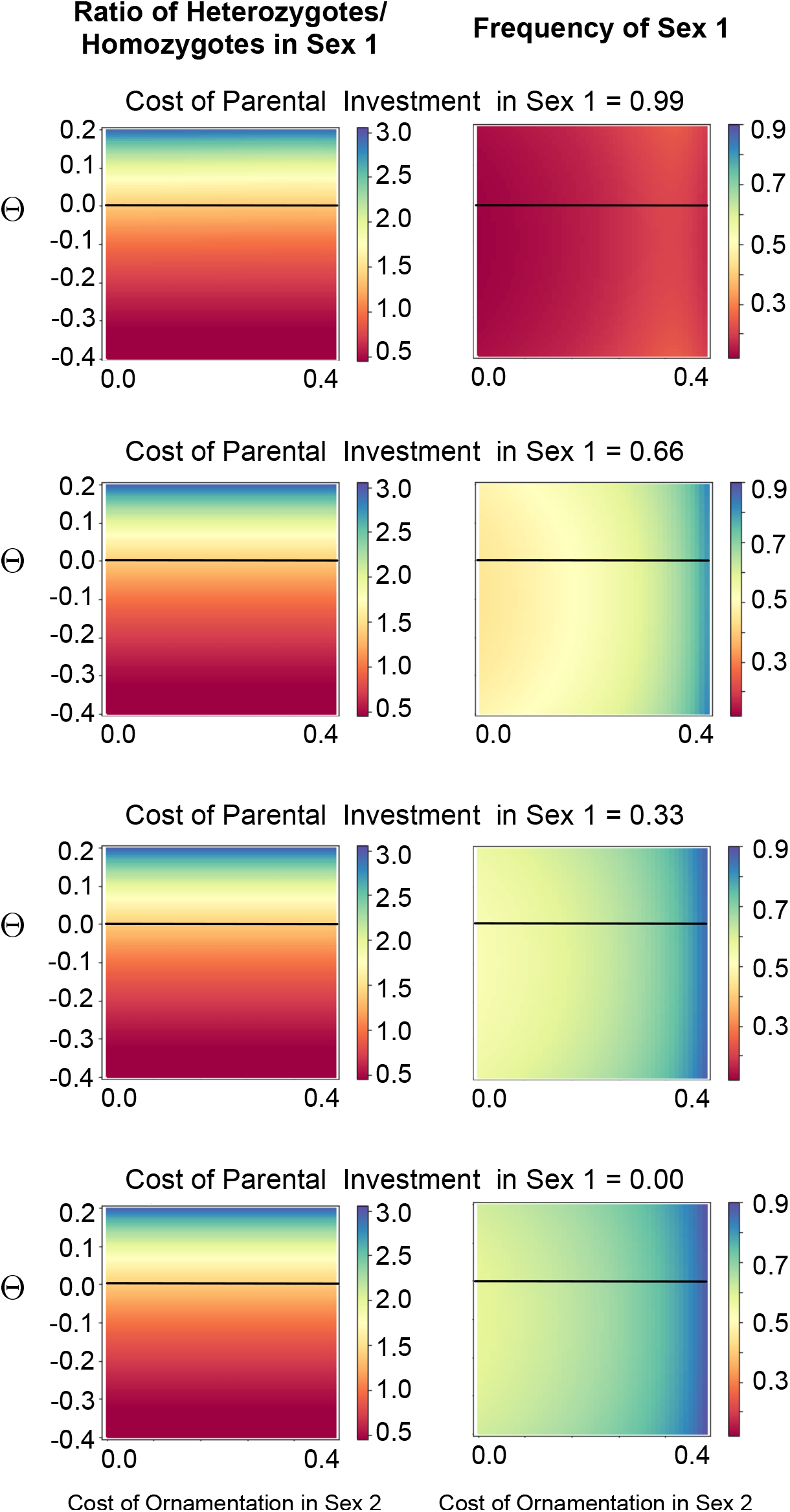
Ratio of Heterozygotes: Homozygotes in Sex 1 and the frequency of Sex 2 for a full range of Θ. The parameter Θ is a measure of the sex difference in immune response through sexual conflict within the MHC as shown in Figure 2. in the main article. It represents the effect of allelic diversity on sex-specific fitness of Sex 1 relative to Sex 2. When Θ = 0, there is so sex-specific difference in immune response. The parameter Θ has an effect on the allele diversity ratio. But there is no effect of ornamentation and parental investment (PI) on this ratio. There is no effect of Θ on the frequency of Sex 1. The cost of ornamentation in Sex 2 increases the frequency of Sex 1 while the cost of PI decreases its frequency.

**Figure S.5:**
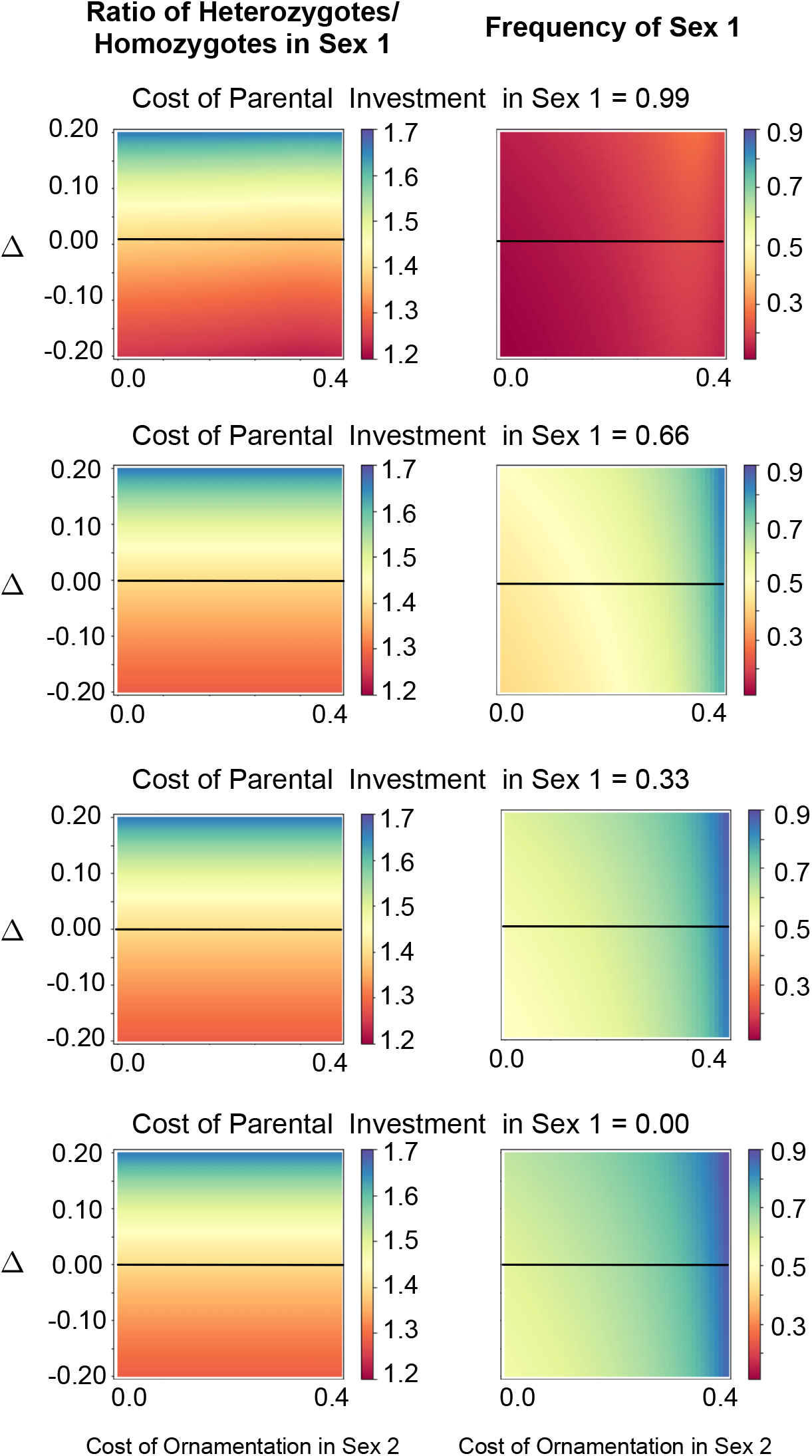
Ratio of Heterozygotes: Homozygotes in Sex 1 and the frequency of Sex 2 for a full range of Δ. The parameter Δ is a measure of the sex difference in immune response through sexual conflict within the MHC as shown in Figure 2. in the main article. It represents the sex-specific fitness effect (that also includes the effect of diversity on sex-specific fitness) of Sex 1 relative to Sex 2. When Δ = 0, there is so sex-specific difference in immune response. There is no effect of ornamentation and parental investment (PI) on the ratio of allele diversity. But Δ has an effect on this ratio. As observed in the previous figure, here too, the cost of ornamentation in Sex 2 and cost of PI in Sex 1 reduce their frequency, respectively while the effect of Δ is not as profound.

## Notes

### Competing Interest Statement

The authors have declared no competing interest.

